# Suppression of amyloid-β secretion from neurons by *cis*-9, *trans*-11-octadecadienoic acid, an isomer of conjugated linoleic acid

**DOI:** 10.1101/2020.09.13.295642

**Authors:** Saori Hata, Kazunori Kikuchi, Kuniyuki Kano, Haruka Saito, Yuriko Sobu, Shoichi Kinoshita, Takashi Saito, Takaomi C. Saido, Yoshitake Sano, Hidenori Taru, Junken Aoki, Hiroto Komano, Taisuke Tomita, Shunji Natori, Toshiharu Suzuki

## Abstract

Conjugated linoleic acid (CLA) comprises several geometric and positional isomers of the parental linoleic acid (LA). Two of the isomers, cis-9, trans-11 CLA (c9,t11 CLA) and trans-10, cis-12 CLA (t10,c12 CLA) exert various biological activities. However, the effect of CLA on generation of neurotoxic amyloid-β (Aβ) protein remains unclear. We found that c9,t11CLA significantly suppressed generation of Aβ in primary cultures of mouse neurons. CLA treatment did not affect the levels of β-site APP-cleaving enzyme 1 (BACE1), a component of active γ-secretase complex presenilin 1 amino-terminal fragment (PS1 NTF), or Aβ protein precursor (APP) in cultured neurons. BACE1 activity in lysate of neurons treated with c9,t11 CLA, but not t10,c12 CLA, decreased slightly, although c9,t11 CLA did not directly affect the activity of recombinant BACE1. Interestingly, localization of BACE1 and APP in early endosomes increased in neurons treated with c9,t11 CLA; concomitantly, the localization of both proteins was reduced in late endosomes, where APP is predominantly cleaved by BACE1. c9,t11 CLA and t10,c12 CLA appeared to be incorporated into membrane phospholipids, as the level of CLA-containing lysophosphatidylcholine (CLA-LPC) increased dramatically in neurons incubated with CLA. Taken together, our findings indicate that accumulation of c9,t11 CLA-LPC, but not t10,c12 CLA-LPC, in neuronal membranes suppresses amyloidogenic cleavage of APP, thereby contributing to preservation of brain neurons by suppressing neurotoxic Aβ production in aged subjects.

## Introduction

Conjugated linoleic acid (CLA) is found in meat products from ruminants. Symbiotic bacteria living in the rumens of these animals generate various CLA isomers by bio-hydrogenating polyunsaturated fatty acids such as linoleic acid (LA). Cis-9, trans-11 CLA (c9,t11 CLA) and trans-10, cis-12 CLA (t10,c12 CLA) are the most common naturally occurring products (reviewed by **den Hartigh, 2019**). CLA exerts a range of biological activities: the intake of CLA is beneficial for cardiovascular health and may also help to prevent cancers and inflammation (reviewed by **Bhattacharya 2006**). The biological activities of these compounds are also observed in the central nervous system; in animal studies, CLA exerts beneficial effects on brain disorders. CLA protects neurons from excitotoxicity caused by excess glutamate (**Hunt 2010**), improves or preserve memory (**Gama 2015; Binyamin 2019**), and prevents age-dependent neurodegeneration (**Monaco 2018**), although it remains unclear to what extent CLA is incorporated into the brain (**Fa 2005**). In many studies, CLA has been used as mixture of isomers; consequently, the detailed biological activities of individual CLA isomers remain unclear.

In Alzheimer’s disease (AD), the most common neurodegenerative disorder in the aged population, the two major pathological hallmarks are senile plaques and neurofibrillary tangles (NFTs) in the brain (**Scheltens 2016**). Senile plaques are formed by extracellular accumulation of their principal component, amyloid-β (Aβ), whereas NTFs are intracellular aggregates of highly phosphorylated tau, a microtubule-associated protein (**Mucke & Selkoe 2012**). Soluble Aβ oligomers are thought to trigger neuronal impairment by attacking synapses prior to the appearance of pathological remarks in brain (**Beniloval 2012**). Therefore, Aβ generation is thought to be a primary cause of AD onset.

Aβ is generated from amyloid-β protein precursor (APP) by serial proteolytic cleavages. APP is a type I membrane protein; in neurons, it contains 695 amino acids (APP695), whereas other tissues express the splicing variants APP770 and APP751 at lower levels (**Iijima et al, 2000**). All APP isoforms are subject to post-translational modifications, including *N*-glycosylation in the ER and *O*-glycosylation and phosphorylation in the Golgi complex. Mature APP (mAPP) with *N*- and *O*-glycosylation is cleaved by secretases in the late secretory pathways, whereas immature APP (imAPP) with *N*-glycosylation alone resides in the ER and *cis*-Golgi compartment (**Suzuki & Nakaya, 2008; Thinakaran & Koo, 2008**).

APP is cleaved by *α*-secretase, primarily on the plasma membrane, to generate the membrane-associated carboxy-terminal fragment CTF*α*; this is termed the non-amyloidogenic pathway. In the amyloidogenic pathway, APP that escapes from cleavage on the plasma membrane is subject to endocytosis and alternative cleavage by β-secretase (BACE1) in endosomes to generate CTFβ (**Cole and Vassar, 2008**). CTFβ, including the entire Aβ sequence, is further cleaved by the γ-secretase complex, yielding Aβ peptides that are secreted into the extracellular milieu. The γ-secretase complex cleaves APP CTF at the *ε*-site to release the APP intracellular domain fragment (AICD) into the cytoplasm. The remaining membrane-associated N-terminal region is subject to further processing by the carboxypeptidase-like activity of the γ-secretase complex to generate various types of Aβ with different carboxyl-terminal γ-cleaved sites, including the strongly neurotoxic and aggregation-prone Aβ42 (**Qi-Takahara et al, 2005; Takami & Funamoto 2012**). Mutations of AD causative genes, including *PSENs* and *APP*, alter Aβ generation (**Mullan et al, 1992; Goate et al, 1991; Forman et al, 1997**), and a protective Icelandic mutation of *APP* decreases β-cleavage and facilitates β’-cleavage of APP, thereby decreasing production of neurotoxic Aβ1-40 and Aβ1-42 (**Jonsson et al, 2012; Kimura et al, 2016**). Accordingly, Aβ generation is considered to be a primary cause of AD.

Although it remains controversial whether Aβ generation is altered in patients with sporadic AD, several lines of evidence support the idea that APP processing or APP secretase activities are altered in these cases (**Yanagida et al, 2009; Kakuda et al, 2012; Kakuda et al, 2020; Hata et al, 2011; Hata et al, 2020**). One cause of changes in APP cleavage and secretase activity is membrane lipid composition. Experimental variation of the proportions of various membrane lipids can alter the C-terminal speciation of Aβ (**Qintero-Monzon et al, 2011; Holmes et al, 2012**), and the cholesterol level in membrane microdomains (detergent-resistant membranes, DRMs) is reduced in AD brains (**Molander-Melin et al, 2005; Hata et al, 2020**). Because the substrates and enzymes related to AD pathobiology are all membrane proteins that are subject to membrane trafficking in neurons (**Suzuki et al., 2006**), lipids and/or fatty acids may play important roles in the metabolism and functions of AD-related membrane proteins, and thus in cognitive functions (**Snowden et al, 2017**).

In this study, we explored the effect of the most common naturally occurring CLA isomers, c9,t11 CLA and t10,c12 CLA, on amyloid-β (Aβ) generation in primary cultured mouse neurons. In order to determine the specific effects of each compound, highly pure samples of each isomer were used.

## Experimental procedures

### Materials

#### Animals and primary cultured neurons and glia

All animal studies were conducted in compliance with the ARRIVE guidelines (approved #13-0096 in the Animal Studies Committee of Hokkaido University). C57BL/6J mice (RRID:IMSR_JAX:000664; CLEA-Japan, Tokyo, Japan) and *App*^*NL- F/NL-F*^ mice (C57BL/6-App<tm3(NL-F)Tcs, RRID:IMSR_RBRC06343) (**Saito et al, 2014**) were housed in a specific pathogen–free (SPF) environment with a microenvironment vent system (Allentown Inc., Allentown, NJ, USA).

Mixed mouse cortical and hippocampal neurons were cultured using a modification of a previous method (**Bartlett and Banker, 1984**). Briefly, the cortex and hippocampus of mice at embryonic day 15.5 were dissected, and neurons were dissociated in a buffer containing papain and cultured at 5 × 10^4^ cells/cm^2^ in Neurobasal Medium (Cat#21103049, Gibco/Thermo Fisher Scientific, Waltham, MA, USA) containing 2% (v/v) B-27 Supplement (Cat#17504044, Gibco/Thermo Fisher Scientific), Glutamax I (4 mM, Cat#35050061, Gibco/Thermo Fisher Scientific), heat-inactivated horse serum (5% v/v, Cat# 26050088, Gibco/Thermo Fisher Scientific), and penicillin–streptomycin (Cat#35050061, Gibco/Thermo Fisher Scientific) on Costar 6-well plates (Cat#3516, Corning, Corning, NY, USA) or Nunc Lab-Tek II Chambered Coverglass (Nalgene Nunc/Thermo Fisher Scientific, Rochester, NY, USA) coated with poly-L-lysine hydrobromide (Cat#P2636, Sigma-Aldrich, St. Louis, MO, USA) (**Chiba et al, 2014**). Neurons cultured for the indicated periods (expressed as days *in vitro*, DIV) were treated with LA (Cat#L1376, Sigma-Aldrich), c9,t11 CLA (Cat# 16413, Sigma-Aldrich), or t10,c12 CLA (Cat#90145, Cayman Chemical, Ann Arbor, MI, USA). In a separate study, neurons were cultured for 24 h with LA, c9,t11 CLA, or t10,c12 CLA (10 μM) in the presence of human Aβ40 (20 nM, Cat#4307-v, Peptide Institute, Osaka, Japan). To quantify exogenous human Aβ40, the levels of Aβ in the medium were assayed by sandwich ELISA (sELISA). Cell viability was evaluated by the MTT assay (Cat #M009, Dojindo Molecular Technologies, Inc., Kumamoto, Japan), and cell toxicity was evaluated using the LDH Cytotoxicity Detection Kit (Cat #MK401; Takara Bio, Shiga, Japan).

### Antibodies, immunoblotting, immunostaining, and Aβ assay

Mouse monoclonal anti-*α*-tubulin (clone 10G10,1:2000, Cat#¥017-25031, Fujifilm-Wako, Osaka, Japan), rabbit monoclonal anti-BACE1 (1: 500, D10E5 Cat#5606, RRID:AB_1903900; Cell Signaling Technologies, Danvers, MA, USA), mouse monoclonal anti-EEA1 (1:500, clone 14, Cat #610457, RRID: AB_397830, BD Biosciences, San Hose, CA, USA), rabbit polyclonal anti-EEA1 (1:500, H-300, Cat#sc-33585, RRID: AB_2277714, Santa Cruz Biotechnology, Dallas, TX, USA), mouse monoclonal anti-Rab7 (1:500, Cat#ab50533, RRID: AB_882241, Abcam, Cambridge, UK), rabbit monoclonal anti-Rab7 (1:500, D95F2, Cat#9367, RRID: AB_1904103, Cell Signaling Technologies), and mouse monoclonal anti-APP (1:500, 22C11; Cat#Mab348, RRID: AB_94882, Merck Millipore/Sigma, Burlington, MA, USA) antibodies were purchased from the indicated suppliers. Rabbit polyclonal anti-APP (1:10,000, 369**; Oishi et al, 1996**) and rabbit polyclonal PS1 NTF (1:10,000, Ab14; **Xu et al, 1998**) antibodies were kindly supplied by Dr. S. Gandy. The amounts of protein were measured using a BCA protein assay kit (Cat#23225, Thermo Fisher Scientific).

Proteins were resolved by Tris-glycine buffered sodium dodecyl sulfate (SDS)– polyacrylamide gel electrophoresis (8% and 15% polyacrylamide gels) and transferred onto nitrocellulose membrane (Cat# P/N66485, Paul Corp., Pensacola, FL, USA) for immunoblotting. Membranes were blocked in 5% non-fat dry milk (barcode 4 902220 354665, Morinaga Milk Industry, Tokyo, Japan) in TBS-T (20 mM Tris-HCl [pH 7.6], 137 mM NaCl, 0.05% Tween 20 [sc-29113, Nacalai Tesque, Kyoto, Japan], probed with the indicated antibodies in TBS-T, and washed with TBS-T. Immunoreactive proteins were detected using Clarity Western ECL substrate (Cat#1705061, Bio-Rad, Hercules, CA, USA) and quantitated on a LAS-4000 (FUJIFILM, Tokyo, Japan). Results are derived from independent experiments, and numbers of experiments (n) are indicated in the figure legends.

Primary neurons from wild-type mice were cultured *in vitro* for 9–12 days (DIV 9-12) and treated with 10 μM LA or CLA for 24 h. Neurons were fixed with 4% paraformaldehyde in PBS, treated with 0.2% Triton X-100 in PBS, and blocked with 1% horse serum and 1% goat serum in PBS. The neurons were incubated with the indicated primary antibodies for 12 h, followed by incubation with Alexa Fluor 488– conjugated donkey anti-mouse IgG (Cat#A-21202, RRID: AB_141607, Invitrogen/Thermo Fisher Scientific), Alexa Fluor 546–conjugated goat anti-mouse IgG (Cat#A-11003, RRID: AB_2534071, Invitrogen/Thermo Fisher Scientific), Alexa Fluor 488–conjugated donkey anti-rabbit IgG (Cat#ab150065, Abcam), and Alexa Fluor 546– conjugated goat anti-rabbit IgG (Cat#A-11010, RRID: AB_2534077, Invitrogen/Thermo Fisher Scientific). Fluorescence images were acquired on a fluorescence microscope (BZ-X710; Keyence, Osaka, Japan), and quantitative analysis and colocalization efficiency were performed with Fiji/ImageJ and the Colco2 Fiji plugin (ImageJ-Fiji-ImgLib; http://Fiji.sc or http://imageJ.net).

Mouse monoclonal anti-Aβ40 (4D1) and Aβ42 (4D8) carboxy-terminal specific antibodies were as described (**Tomita et al, 1998**). The Fab’ fragment of rabbit polyclonal anti-mouse Aβ(1-16) IgG conjugated with horseradish peroxidase (Cat#27720, IBL, Fujioka, Japan) was used for the detection of mouse Aβ40. Biotinylated mouse monoclonal anti-human Aβ(1–16) IgG 82E1 was purchased from IBL (Cat#10326, RRID: AB_1630806) and used for detection of human Aβ. Mouse Aβ40 was quantified by sELISA consisting of 4D1 and anti-mouse Aβ(1–16), and human Aβ42 was quantified by sELISA consisting of 4D8 and 82E1 (**Mizumaru et al, 2009**).

#### In vitro β-secretase assay

BACE1 activity was examined using the β-secretase activity assay kit (Abcam, Cat. #ab65357). In short, neurons were homogenized with BACE1 extraction buffer (contents are not opened), and the lysate (100 μg protein) or 4 μg of recombinant human BACE1 (Cat. # 931-AS; R&D Systems, Minneapolis, MN) in extraction buffer was incubated at 37°C for 1 h with a secretase-specific peptide conjugated to two reporter molecules. After the reaction was stopped, Ex/Em (335 nm/495 nm) was measured.

### LC-MS/MS analysis

Neurons (DIV 10, ∼5 × 10^5^ cells) derived from wild-type mice were treated with 10 μM LA or CLA for 24 h. Lipids were recovered with acidic methanol (pH 4.0, 200 µL) that contained 1 µM 17:0-LPC as an internal standard and stored at −80°C. LC-MS/MS analysis was performed as described previously (**Kano et al, 2019**) with some modifications. Briefly, LC was performed using a C18 column (CAPCELL PAK C18 (1.5 mm I.D. × 250 mm; particle size, 3 µm) with gradient elution using solvent A (5 mM ammonium formate in 95% (v/v) water, pH 4.0) and solvent B (5 mM ammonium formate in 95% (v/v) acetonitrile, pH 4.0) at 150 µL/min. Gradient conditions were as follows: hold 50% B for 0.5 min, linear gradient to 100% B over 18 min, hold 100% B for 7 min, and then return to the initial condition. MS analysis was performed by selective reaction monitoring (SRM) in positive mode. LA- and CLA-LPC SRM transitions were as follows: Q1 *m/z* 520.34, Q3 *m/z* 184.0.

### Statistical analysis

Data are expressed as means ± SE. Statistical differences were assessed using Student’s t-test or one-way ANOVA, combined with the Tukey–Kramer post-hoc test and Dunnett’s test for multiple comparisons (GraphPad Prism 8 software). A *p* value of <0.05 was considered significant. No sample size calculation or tests for normal distribution or outliers were performed. The study was not pre-registered.

## Results

### Suppression of Aβ generation in neurons treated with c9,t11 CLA

To explore the effect of CLA on Aβ generation, we treated mixed cortical and hippocampal neurons (DIV 14–17) from human APP knock-in mice (*App*^*NL-F/NL-F*^) with 10 μM LA, c9,t11 CLA, or t10,c12 CLA for 48 h, and then quantified humanized Aβ42 secreted into medium by sELISA (**Fig. 1A**). Interestingly, c9,t11 CLA decreased Aβ42 generation significantly relative to LA, but t10,c12 CLA had no effect on Aβ42 generation. This observation shows that c9,t11 CLA, but not t10,c12 CLA, has the ability to suppress Aβ generation We did not quantify humanized Aβ40 because the APP-KI mouse predominantly generates Aβ42 (**Saito et al, 2014**). To confirm the effect of c9,t11 CLA on Aβ generation, we treated mixed cortical and hippocampal neurons (DIV7) from wild-type (WT) mice for 24 h with 10 μM LA, c9,t11 CLA, or t10,c12 CLA, and then assessed endogenous mouse Aβ40 secreted from neurons (**Fig. 1B**). Generation of mouse Aβ40 was significantly lower in neurons treated with c9,t11 CLA, but not t10,c12 CLA, than in neurons treated with LA, indicating that the suppression of Aβ generation by c9,t11 CLA was reproducible. Levels of endogenous mouse Aβ42 were too low to quantify using cultured neurons.

**Figure 1.**
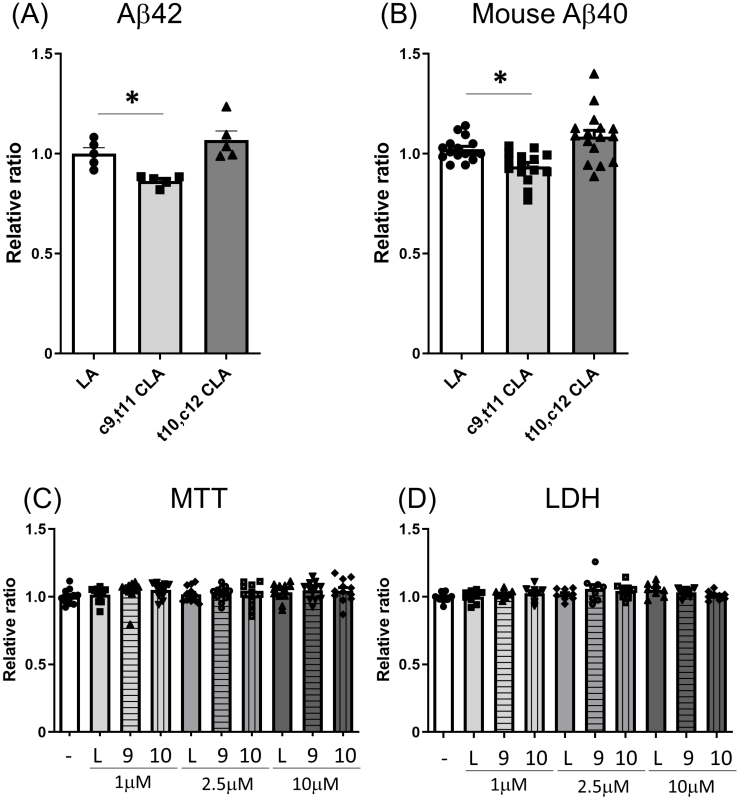
Aβ generation and viability of neurons treated with LA and CLA. (**A, B**) Aβ peptides in medium generated from primary cultured neurons derived from hippocampus and cortex of APP -KI (**A**) and wild-type (**B**) mouse embryos. (**A**) Neurons of APP-KI (*AppNL-F/NL-F*) mice were cultured for 14 to 17 days (DIV 14–17). Neurons were cultured for an additional 48 h in fresh medium containing 10 μM LA, c9,t11 CLA, or t10, c12 CLA, and human Aβ42 in the medium was quantified. (**B**) Neurons of wild-type mice were cultured for 7 days (DIV 7). Medium was changed as described above, and endogenous mouse Aβ40 in the medium was quantified. Aβ levels are indicated as ratios relative to the level of Aβ generated by neurons treated with LA, which was assigned a reference value of 1.0. (**C, D**) Cell viability and toxicity of neurons. Neurons of wild-type mice were cultured for 12 days (DIV 12). Neurons were further cultured for 48 h in fresh medium containing the indicated concentrations of LA, c9,t11 CLA, or t10, c12 CLA, and cell viability (**C**) and cell toxicity (**D**) were examined. Statistical analysis was performed using Dunnett’s multiple comparison test. P values are provided for comparison with neurons treated with LA (means ± S.E., n=2–3 × 2 times (**A**), n=4 × 4 times (**B**); *, p<0.05), or with vehicle (-) (means ± S.E., n=6 × 2 times (**C, D**))

We next assessed the viability of neurons after treatment with CLA and LA. To this end, we treated neurons (DIV 12) from wild-type mice with the indicated concentration of LA, c9,t11 CLA, t10,c12 CLA, and then examined cell viability by MTT assay (**Fig. 1C**). There were no significant differences among neurons treated with CLA and LA at any concentration. In addition, we measured the levels of lactate dehydrogenase (LDH), an indicator of cytotoxicity and plasma membrane injury, in the medium of mouse primary neurons cultured in the presence of LA and CLA. Again, we detected no significant differences among the conditions (**Fig. 1D**). These data indicate that treatment of neurons with LA and CLA does not affect cell viability, and that the decrease in Aβ generation in neurons treated with c9,t11 CLA was not due to changes in cell viability.

### c9,t11CLA does not promote Aβ degradation in medium of primary cultured neurons and glial cells

Extracellular Aβ is subject to clearance by cellular uptake and proteolytic degradation (**Saido and Leissring, 2012**). The lower levels of Aβ in the medium of neurons treated with c9,t11 CLA could have been the result of faster clearance of Aβ from the medium. To explore this possibility, we investigated whether Aβ clearance or degradation was facilitated by CLA treatment of neurons and glial cells. For this purpose, we treated primary cultured neurons (DIV10–12) and glia of WT mice with or without LA, c9,t11 CLA, and t10,c12 CLA for 24 or 48 h in the presence of human Aβ. Media were collected at the indicated times, and Aβ contents were examined by sELISA or immunoblotting. In media from neurons (**Fig. 2**) and glia (**Supplementary Fig. S1A, B**), Aβ levels were slightly lower than in cell-free media alone. However, the magnitudes of the decrease were almost identical between cells treated with LA and CLA. As in the case of neurons, the viability of glia was not affected by treatment with LA or CLA (**Supplementary Fig. S1C**). Although primary cultured neurons may include small numbers of glial cells, these observations indicates that the reduced level of Aβ in medium of neurons treated with c9,t11 CLA was not due to Aβ clearance or degradation by neurons and glia, which secrete Aβ degrading enzymes (**Kidana et al, 2018**).

**Figure 2.**
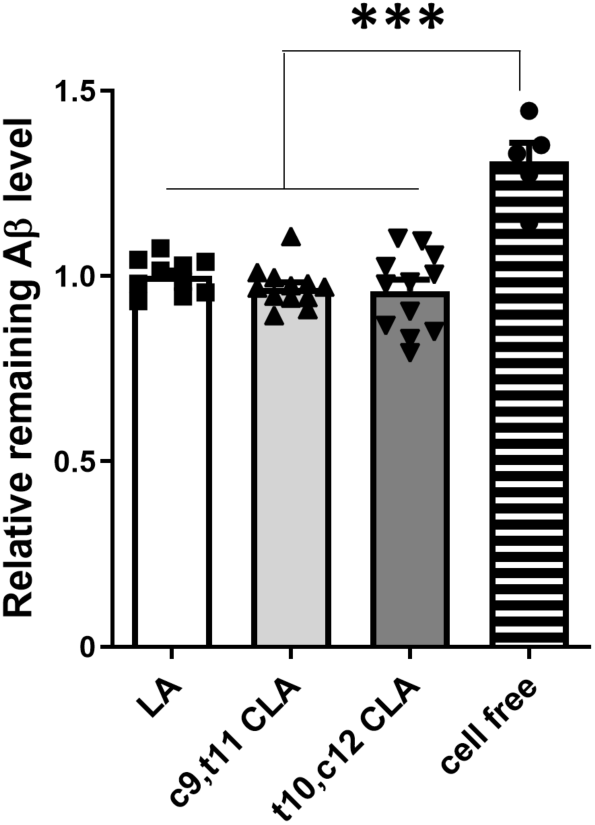
Aβ clearance in medium of neurons treated with LA and CLA. Neurons of wild-type mice were cultured for 10 to 12 days (DIV10–12), and further cultured in fresh medium containing 10 μM LA, c9,t11 CLA, and t10, c12 CLA with Aβ42 (10 nM) for 24 h. ‘Cell-free’ indicates medium containing Aβ42 in the absence of neurons. Levels of Aβ42 in medium were quantified by sELISA, and are indicated as ratios relative to the level of Aβ in medium of neurons treated with LA, which was assigned a reference value 1.0. No significant differences were observed among LA, c9,t11 CLA, and t10, c12 CLA (Tukey’s multiple comparison test). P values (***, p<0.001) are shown for comparison between LA, c9,t11 CLA, or t10,c12 CLA and cell-free (means ± S.E., n=5 × 2 times).

### c9,t11 CLA decreases BACE1 activity in neuronal lysate, but not *in vitro*

To investigate how neurons treated with c9,t11 CLA decrease Aβ generation, we first assessed the level of β-site APP-cleaving enzyme (BACE1), which is the enzyme primarily responsible for amyloidogenic cleavage of APP (**Vassar et al, 1999; Kimura et al, 2016**). Because CLA alters expression of BACE1 but not ADAM10/*α*-secretase in SH-SY5Y cells (**Li et al, 2011**), we examined protein expression of APP, BACE1, and presenilin 1 amino terminal fragment (PS1 NTF), a catabolic component of active γ-secretase complex, in neurons treated with CLA and LA. The levels of these proteins, as well as *α*-tubulin (used as a loading control), were measured by immunoblotting with the corresponding antibodies (**Fig. 3A**). Cellular levels of APP, BACE1, and PS1 NTF were almost identical among neurons treated with LA, c9,t11 CLA, and t10,c12 CLA, and no significant differences were observed (**Fig. 3B–D**). These results suggest that differences in the levels of substrate APP and APP-cleaving enzymes are not the primary cause of the reduction in Aβ generation in neurons treated with c9,t11 CLA.

**Figure 3.**
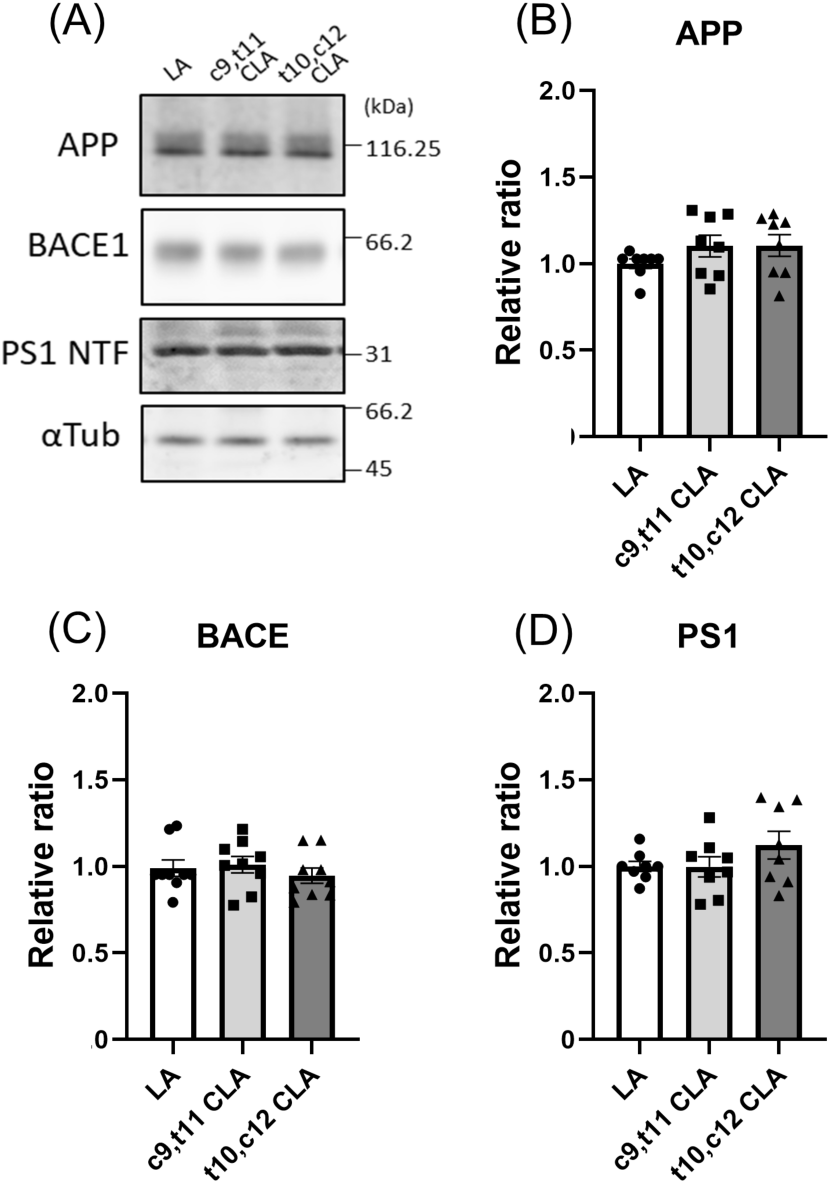
Protein expression of neurons treated with LA and CLA. Immunoblotting **(A)** and quantification **(B– D)** of protein levels of APP, BACE1, PS1 NTF, and *α*-tubulin in neurons treated with LA, c9,t11 CLA, and t10,c12 CLA. Primary cultured neurons derived from hippocampus and cortex of wild-type mouse embryos (DIV 10-12) were cultured for 24 h with 10 μM LA, c9,t11 CLA, and t10,c12 CLA. These neurons were lysed, and the lysate (15 μg protein) was analyzed by immunoblotting with anti-APP, BACE1, PS1 NTF, and *α*-tubulin antibodies. The numbers in panel A indicate molecular size (kDa). The densities of each protein band were quantified and normalized against the density of *α*-tubulin. The levels of protein in cells treated with CLA were compared with the level in cells treated with LA, which was assigned a reference value of 1.0. Statistical analysis was performed using Dunnett’s multiple comparison test; no significant difference was detected (means ± S.E., n=7).

Next, we examined BACE1 activity. Specifically, we assayed lysates of wild-type mouse neurons treated with CLA and LA for BACE1 activity *in vitro*. We treated primary cultured neurons (DIV 9–12) from wild-type mice with 10 μM LA, c9,t11 CLA, or t10,c12 CLA for 24 h and assayed their lysates for BACE1 activity (**Fig. 4A**). BACE1 protein level was not affected by treatment with CLA or LA (**Fig. 3**), but lysates prepared with this assay system revealed that BACE1 enzyme activity was slightly but significantly reduced in neurons treated with c9,t11 CLA, but not t10,c12 CLA, relative to those treated with LA (**Fig. 4A**).

**Figure 4.**
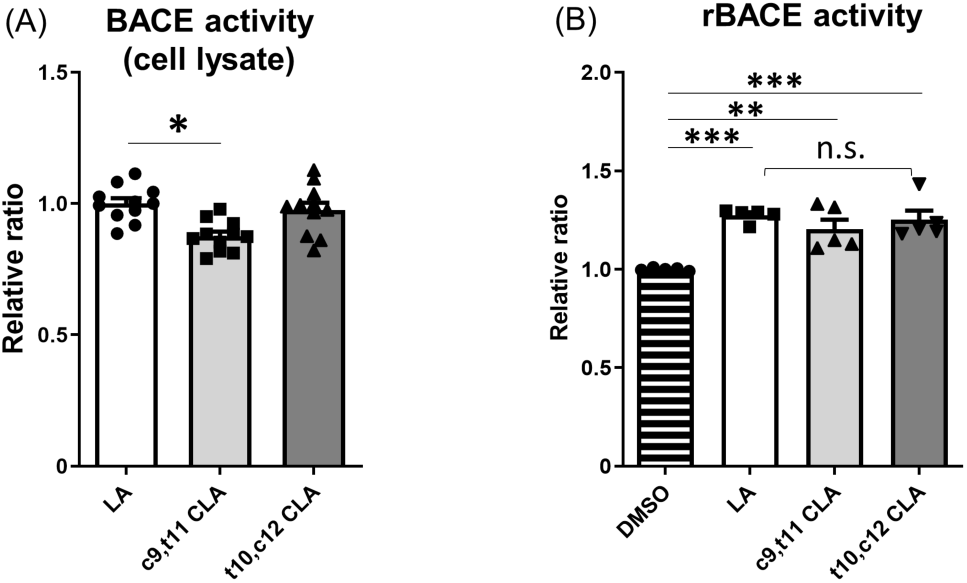
BACE1 activity in lysate of neurons treated with LA and CLA, and activity of recombinant human BACE1 in the presence of LA and CLA. (**A**) Primary cultured neurons derived from hippocampus and cortex of wild-type mouse embryos (DIV 9–12) was cultured for 24 h with 10 μM LA, c9,t11 CLA, or t10,c12 CLA. BACE1 activity in lysates (100 μg protein) of neurons was examined, and activities of neurons treated with CLA are indicated as ratios relative to the activity of neurons treated with LA, which was assigned a reference value 1.0. Statistical analysis was performed using Dunnett’s multiple comparison test. P values are provided for the comparison with neurons treated with LA (means ± S.E., n=2–4 × 4 times; *, p<0.05). (**B**) Human recombinant BACE1 (rBACE1) was assayed in the presence or absence of 10 μM LA, c9,t11 CLA, or t10,c12 CLA. The activity of rBACE1 in the absence of LA and CLA (DMSO) was assigned a reference value of 1.0, and the relative activities of rBACE1 in the presence of LA and CLA are indicated as relative ratios. Statistical analysis was performed using Tukey’s multiple comparison test. P values are provided for the comparison with rBACE1 without LA and CLA (means ± S.E., n=1-2 × 3times; **, p<0.01; ***, p<0.005).

We next investigated whether c9,t11 CLA directly affects BACE1 activity *in vitro* (**Fig. 4B**). To this end, we incubated recombinant BACE1 with or without LA, c9,t11 CLA, and t10,c12 CLA, and then assayed enzymatic activity following addition of the BACE1 substrate. In the presence of LA or CLA, BACE1 activity was slightly higher than in the vehicle (DMSO) control. However, we observed no remarkable differences in BACE1 activity among enzyme solutions containing LA, c9,t11 CLA, and t10,c12 CLA (**Fig. 4B**). These results clearly show that c9,t11 CLA does not directly and specifically activate BACE1. The slight increase of BACE1 activity in the presence of LA and CLA may be due to stabilization of BACE1, which is a membrane protein and could therefore be stabilized in the presence of lipids.

It is difficult to reasonably interpret these results. The differences in BACE1 activity in the neuronal lysates used in this study may reflect alterations of membrane lipid composition or intracellular localization of BACE1 following treatment of neurons with c9,t11 CLA. Therefore, we next explored the localization of BACE1 and APP in neurons, and also analyzed the membrane lipid compositions of neurons.

### Reduced localization of APP with BACE1 in late endosomes of neurons treated with c9,t11 CLA

APP cleavage by BACE1 occurs in detergent-resistant membrane domains (DRMs) or lipid rafts (**Cordy et al, 2003; Puglielli et al, 2003; Kalvodova et al, 2005; Saito et al, 2008**) of acidic organelles following plasma membrane endocytosis of APP (**Cole and Vassar, 2008**). Hence, we first examined the localization of APP and BACE1 in neurons with or without CLA or LA treatment, and compared the colocalization efficiency of APP with BACE1 among the samples (**Fig. 5**). Colocalization of APP with BACE1 in neurons is shown (**Fig. 5A, first row**). Moderate colocalization with a Pearson’s R value of ∼0.5 was observed in neurons treated with LA (a value of 1.0 indicates perfect colocalization, 0 indicates random localization, and −1 indicates perfect exclusion).

**Figure 5.**
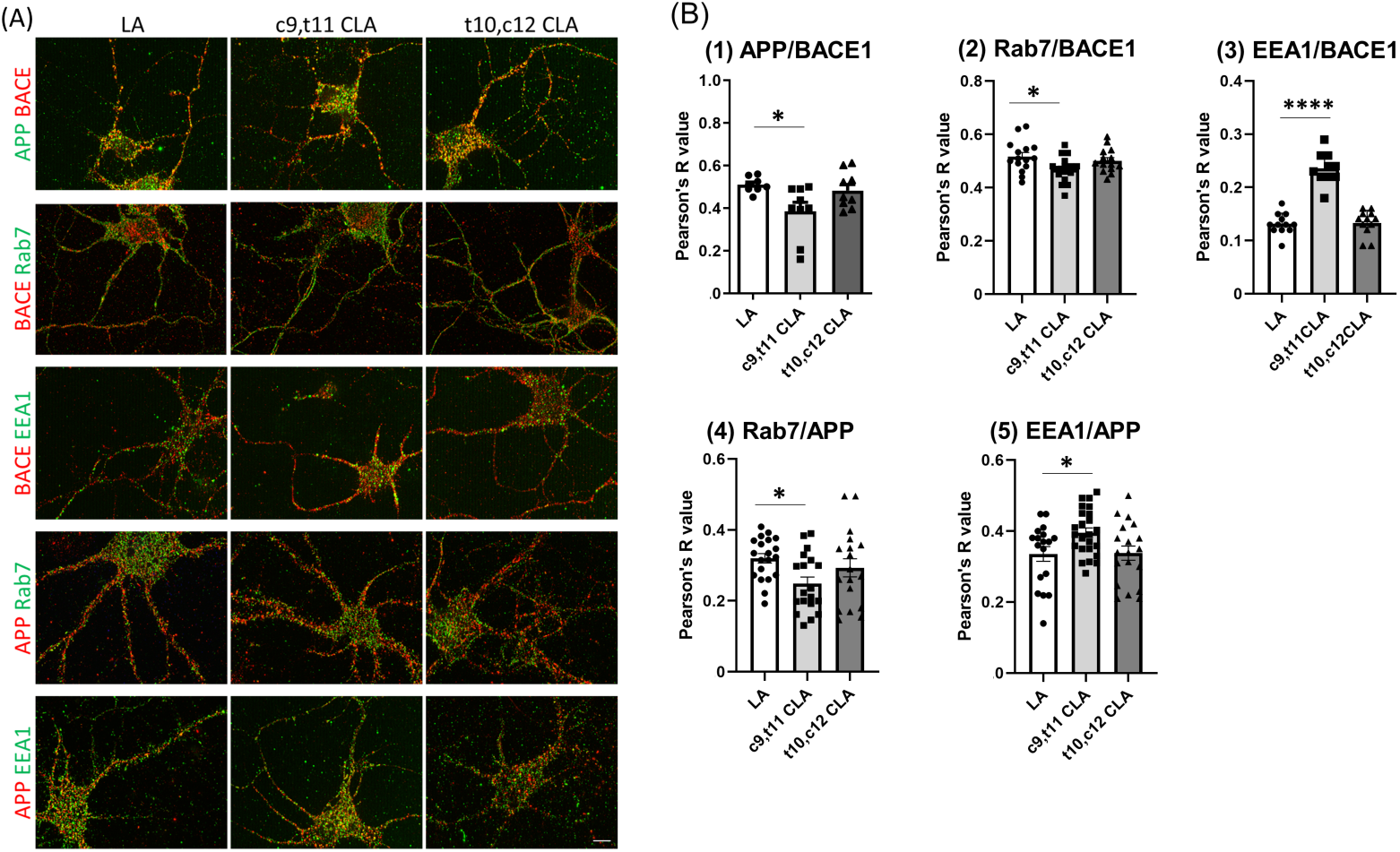
Colocalization of BACE1 and APP with Rab7 and EEA1 in neurons treated with LA and CLA. **(A) Immunostaining**. Primary cultured neurons derived from hippocampus and cortex of wild-type mouse embryos (DIV 10–12) were cultured for 24 h with 10 μM LA, c9,t11 CLA, or t10,c12 CLA, and subject to immunostaining with a combination of the indicated antibodies. First row: mouse anti-APP (green) and rabbit anti-BACE1 (red) antibodies. Second row, rabbit anti-BACE1 (red) and mouse anti-Rab7 (green) antibodies. Third row, rabbit anti-BACE1 (red) and mouse anti-EEA1 (green) antibodies. Fourth row, rabbit anti-APP (red) and mouse anti Rab7 (green) antibodies. Fifth row, rabbit anti-APP (red) and mouse anti-EEA1 (green) antibodies. Scale bar, 10 μm. **(B)** Colocalization efficiency. Colocalization efficiency is indicated by Pearson’s R value. (1) Colocalization of BACE1 with APP. (2) Colocalization of BACE1 with Rab7. (3) Colocalization of BACE1 with EEA1. (4) Colocalization of APP with Rab7. (5) Colocalization of APP with EEA1. Statistical analysis was performed using Dunnett’s multiple comparison test. P values are provided for the comparison with LA treatment (means ± S.E., n=5–7 × 2 times; *, p<0.05; ****, p<0.0001).

However, Pearson’s R value was significantly lower in neurons treated with c9,t11 CLA than in those treated with LA, whereas t10,c12 CLA had no effect (**Fig. 5B (1)**). The observation suggests that colocalization of BACE1 with APP decreased in neurons treated with c9,t11 CLA.

We next examined colocalization of BACE1 with Rab7, a late endosomal marker (**Fig. 5A, second row**). Interestingly, colocalization between BACE1 and Rab7 significantly decreased in neurons treated with c9,t11 CLA, but not t10,c12 CLA or LA (**Fig. 5B (2)**), suggesting less localization of BACE1 in late endosomes, where BACE1 cleaves APP in an acidic milieu. We also examined colocalization of BACE1 with EEA1, an early endosomal marker (**Fig. 5A, third row**). In contrast to Rab7, colocalization of BACE1 with EEA1 remarkably increased in neurons treated with c9,t11 CLA relative to those treated with t10,c12 CLA, or LA (**Fig, 5B (3)**). These observations suggest that membrane transport of BACE1 into late endosomes from early endosomes may be delayed, and that BACE1 tends to remain in early endosomes, resulting in weaker BACE1activity.

We also analyzed the localization of APP with Rab7 and EEA1 (**Fig. 5A, fourth and fifth rows**). Colocalization of APP with Rab7 decreased significantly in neurons treated with c9,t11 CLA, whereas colocalization of APP with EEA1 was greater in neurons treated with c9,t11 CLA than in those treated with t10,c12 CLA or LA (**Fig, 5B (4) and (5)**). These alterations were identical to those observed for BACE1. Overall, these findings imply that the decrease in colocalization of APP with BACE1 in late endosomes, where BACE1 actively cleaves APP, causes the reduction in Aβ generation in neurons treated with c9,t11 CLA.

### Altered phospholipid composition of neurons treated with CLA

The regulation of the transition from early to late endosomes and membrane protein trafficking into the late endosome from the early endosome is controversial, and the details of how APP and BACE1 are regulated remain unknown. In general, membrane lipid composition affects the localization and function of membrane proteins. Therefore, we used a reverse-phase LC/MS system to analyze phosphatidylcholine (PC), a major lipid constituent of the membrane, in neurons treated with LA or CLA. In neurons cultured in the presence of LA (18:2) or CLA (18:2), the peak of the corresponding PC species (*i.e*. 18:2-containing PC species) increased (data not shown). However, we could not determine the level of PC-containing CLA because CLA- and LA-containing PC could not be satisfactorily separated by our reverse-phase LC (data not shown). Therefore, we focused our analysis on lysophosphatidylcholine (LPC), which has only one acyl group at the *sn*-1 or *sn*-2 position of the glycerol backbone. Because LPC has only one acyl chain, we expected that LA- and CLA-containing LPC could be separated using our current LC methods. In control DMSO (vehicle)-treated neurons, we observed two peaks of LA-LPC corresponding to 1-LA-LPC and 2-LA-LPC (Okudaira et al, 2014) (**Fig. 6A**). The levels of both 1-LA-LPC and 2-LA-LPC were significantly higher in cells treated with LA than in those treated with control DMSO (**Fig. 6B**), suggesting that exogenous LA is incorporated into LPC.

**Figure 6.**
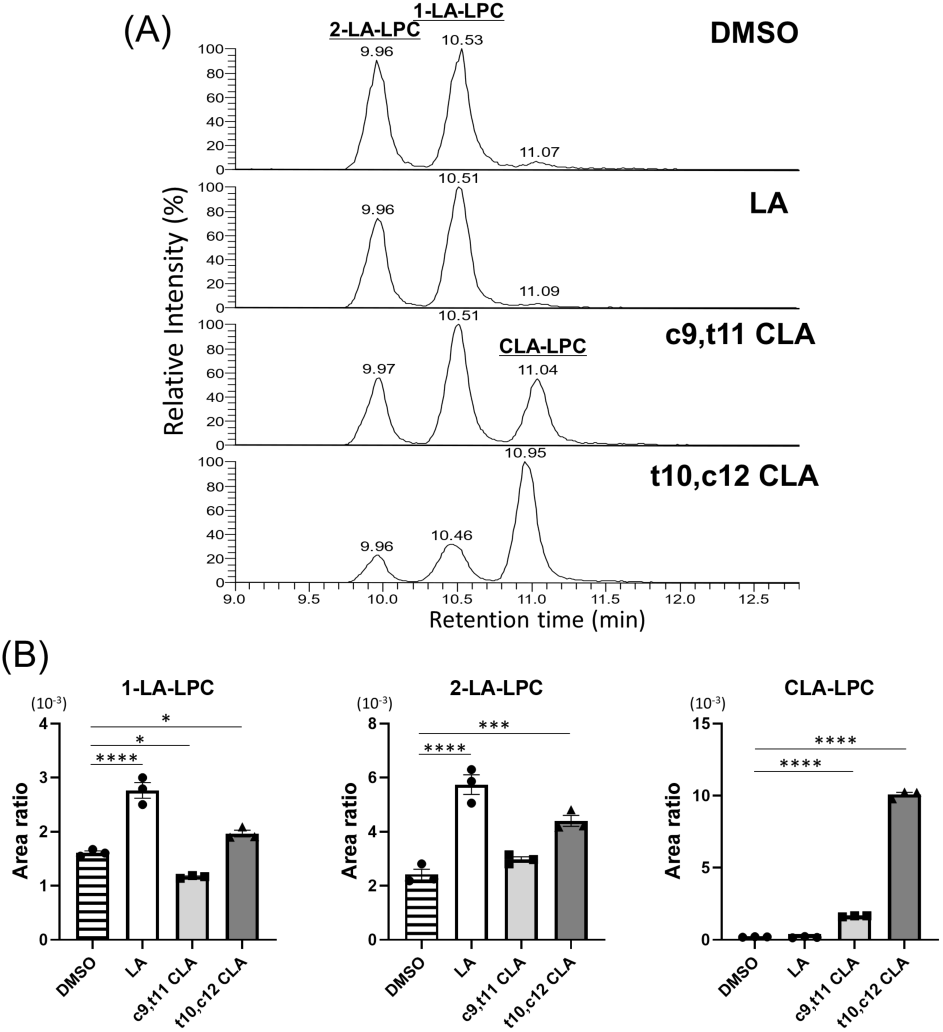
LC-MS/MS analysis of LPC in neurons treated with LA and CLA. **(A)** Representative elution profiles of LA-LPC and CLA-LPC. LPC from neurons treated with 10 μM of linoleic acid (LA), c9,t11 CLA, and t10,c12 CLA, or with vehicle alone (DMSO) was analyzed by LC-MS/MS. **(B)**Comparison of LA-LPC and CLA-LPC levels in neurons treated with LA and CLA. Peak areas are calculated relative to the area of the internal standard, 17:0-LPC (1 μM), and indicated as area ratios (1.0 for 17:0-LPC). The area ratio was compared to that of control neurons (DMSO). Statistical analysis was performed using Dunnett’s multiple comparison test. P values are provided for the comparison with DMSO control (means ± S.E., n= 3; *, p<0.05; **, p<0.01; ***, p<0.001, ****, p<0.0001).

In cells treated with c9,t11 CLA or c10,t12 CLA, we detected an extra peak with a delayed retention time **(Fig. 6A**). The peak appearing in neurons treated with c9,t11 CLA had a retention time slightly different from that of the peak in neurons treated with t10,c12 CLA in several previous studies, although the two peaks were not separated clearly in our LC system. The retention time of CLA on reverse-phase LC is longer than that of LA (**Banni et al, 1994**), suggesting that these extra peaks correspond to CLA-LPCs. The peak in c9,t11 CLA-treated neurons was assigned as c9,t11CLA-LPC, and the peak observed in t10, c12 CLA-treated neurons as t10,c12 CLA-LPC.

We compared the levels of LA-LPC and CLA-LPC in neurons treated with LA or CLA to those in control neurons as an area ratio relative to 17:0-LPC (**Fig. 6B**). Again, the levels of both 1-LA-LPC and 2-LA-LPC were dramatically increased by treatment with LA, whereas the level of 1-LA-LPC decreased modestly following treatment with c9,t11 CLA, and the level of 2-LA-LPC increased modestly following treatment with t10,c12 CLA. The levels of CLA-LPC(s) were greatly increased by treatment with either c9,t11 CLA or t10,c12 CLA. These results suggest that, as with LA, CLA is also incorporated into membrane phospholipids including PC. Because LPCs make up 0.1– 1% of PCs in many cell types, including neurons, it is likely that some CLA-PC was converted to CLA-LPC. Thus, phospholipids containing c9,t11 CLA, but not t10,c12 CLA, may change membrane lipid composition, leading to reduction in Aβ generation by regulating the localization and function of APP and BACE1.

## Discussion

Conjugated linoleic acid (CLA), which comprises various positional and stereoisomers of octadecadienoic acid, exerts multiple bioactive functions. Since the discovery of its anticarcinogenic effects (**Pariza and Hargraves 1985**), subsequent studies revealed additional biological activities that suppress adiposity (**den Hartigh et al, 2013**), atherosclerosis (**Bruen et al, 2019**), diabetes (**Clement et al, 2002**), and inflammation (**Viladomiu et al, 2016**) (reviewed in **Bhattacharya et al, 2006; den Hartigh 2019**). In many of these studies, CLA was used *in vivo* or *in vitro* as an isomeric mixture consisting largely of c9,t11 CLA and t10,c12 CLA. The activities of CLA are also beneficial to the central nervous system and in neurons. A mixture of the two major CLA isoforms, c9,t11 CLA and t10,c12 CLA, is incorporated into rat brain following a single-dose gavage (**Fa et al, 2005**). In addition, a diet containing CLA increases phospholipase A2 expression and activity in rat hippocampus, which correlates with memory enhancement (**Gama et al, 2015**). Furthermore, in a mouse model of neuropsychiatric lupus, CLA supplementation suppressed age-associated neuronal damage *in vivo* (**Monaco et al, 2018**), and c9,t11 CLA increased the proliferation of neural progenitor cells *in vitro* (**Wang et al, 2011**). Administration of punicic acid, from which CLA is generated as a metabolite, preserves memory and decreases Aβ accumulation in a mouse model of AD (**Binyamin et al, 2019**). Together, this evidence supports the idea that CLA can prevent neurodegenerative disease in aged populations. The most common age-associated neurodegenerative disease is AD, and the neurotoxicity of Aβ oligomers is believed to trigger the impairment of neurons. However, the evidence supporting a role for CLA in AD prevention and care is limited, and it remains unclear how CLA prevents neurons in AD pathologies and which isomer plays an important role in neuronal function for AD pathobiology. Therefore, in this study, we focused our analysis on the influence of CLA on Aβ generation in neurons.

Our major finding was that c9,t11 CLA suppressed Aβ generation, and that this effect was specific for c9,t11 CLA but not t10,c12 CLA. The activity of c9,t11 CLA was confirmed for two Aβ species in two types of mouse primary cultured neurons: endogenous mouse Aβ40 in neurons of wild-type mice and humanized Aβ42 in human (*App*^*NL-F/NL-F*^) APP-KI mice (**Saito et al, 2014**). c9,t11 CLA did not affect the viability of mouse neurons. Because CLA exerts multiple biological activities in cells and individual subjects (reviewed by **den Hartigh, 2019**), the reduction in Aβ generation in neurons mediated by c9,t11 CLA may be the consequence of various biological functions. Among them, we observed that the opportunity of APP to interact with BACE1 in the late endosome was reduced in neurons treated with c9,t11 CLA. This was due to reduced localization of BACE1 and APP into late endosomes, where APP is subject to amyloidogenic cleavage by BACE1, along with alteration of membrane phospholipid composition. Our study provides the first evidence that CLA is incorporated into LPC in neuronal membranes. Although structural analyses of the compounds have not yet been completed, c9,t11 CLA-LPC and t10,c12CLA-LPC are potentially generated as is LA-LPC. Therefore, a membrane lipid composition modified with c9,t11 CLA-LPC, but not t10,c12 CLA-LPC, may contribute to suppression of amyloidogenic processing of APP. Because CLA can be incorporated in many other lipids in the cells, including phospholipids, tri-, di-, and mono-acyl glycerols, and cholesteryl esters, we cannot rule out the possibility that other lipid components in the membrane may be effective for suppressing Aβ generation.

In a separate study in which an AD mouse model was fed c9,t11 CLA, c9,t11 CLA-LPC was detected in brain membranes, concomitant with a reduction in amyloid plaques (**Fujita et al, submitted elsewhere**). It remains unclear whether the biological effects of c9,t11 CLA observed *in vivo* are due to the same effect observed in cultured neurons in this study. Nonetheless, taken together with Fujita’s report, our results indicate that intake of high-purity c9,t11 CLA protects neurons in aged subjects from injury by toxic Aβ. Another major isoform, t10,c12 CLA, has a remarkable ability to decrease adiposity (**Park et al, 1999; den Hartigh et al, 2013**), but a CLA mixture containing t10,c12 CLA induces fatty liver (**Clement et al, 2002; Pang et al, 2019**). By contrast, induction of fatty liver and reduction of body fat were not observed in mice fed diets containing highly pure c9,t11 CLA (**Sano et al**., **unpublished observation**). Therefore, administration of a CLA mixture containing both c9,t11 CLA and t10,c12 CLA to aged subjects may not be beneficial for long-term health. Intake of diets and/or supplements including highly purified c9,t11 CLA, or at least limited amounts of t10,c12 CLA alone, may be effective for preventing neurodegenerative disease such as AD without peripheral side effects.

## Abbreviations used

APP: amyloid-β protein precursor
AD: Alzheimer’s disease
CLA: conjugated linoleic acid
LA: linoleic acid
BACE1: β-site APP-cleaving enzyme 1
NMDA: N-methyl-D-aspartate
AMPA: *α*-amino-3-hydroxy-5-methyl-4-isoxazo-leproionic acid
MTT: 3-(4,5-di-methylthiazol-2-yl)-2,5-diphenyltetrazolium bromide, yellow tetrazole
LDH: lactate dehydrogenase
ELISA: enzyme-linked immunosorbent assay
LPC: lysophosphatidylcholine

---

**Supplementary Figure S1.**
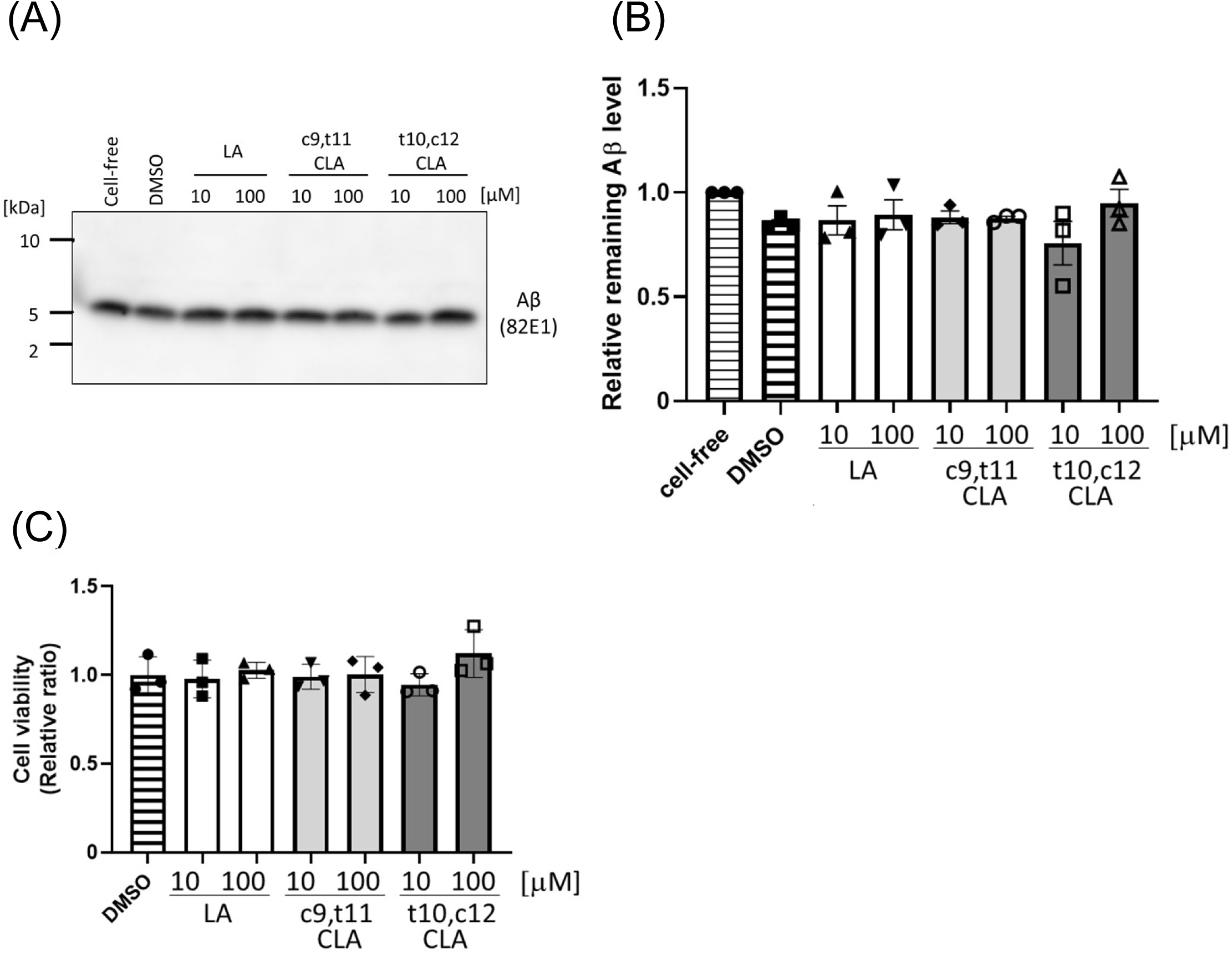
Aβ clearance in medium of glia treated with LA and CLA. Primary cultured glial cells were prepared from cortex of 1–2-day post-natal mouse brains and cultured for 15 days. Glial cells (largely astrocytes) of wild-type mice were cultured for 48 h in fresh medium containing 10 or 100 μM LA, c9,t11 CLA, or t10,c12 CLA with 20 nM Aβ40. **(A, B)** Aβ in medium was quantified by immunoblotting with anti-human Ab antibody (82E1). Band densities are indicated as ratios relative to the level of Aβ in medium in the absence of cells, which was assigned a reference value 1.0. No significant difference was observed among LA, c9,t11 CLA, and t10, c12 CLA using Dunnett’s multiple comparison test (n=3), although Aβ levels tend to be lower in the presence of cells than in cell-free medium (1.0). Numbers (panel A, left side) indicate molecular size (kDa). **(C)**Cell viability was assayed by staining with Alamar blue (Cat. #DAL1025, Thermo Fisher Scientific, Waltham, MA. USA), and is indicated as the ratio relative to the level of cells without treatment (DMSO), which was assigned a reference value of 1.0. No significant difference was observed among LA, c9,t11 CLA, and t10,c12 CLA (Dunnett’s multiple comparison test; n=3).

